# Mitonuclear interactions impact aerobic metabolism in hybrids and may explain mitonuclear discordance in young, naturally hybridizing bird lineages

**DOI:** 10.1101/2023.11.19.567741

**Authors:** Callum S. McDiarmid, Daniel M. Hooper, Antoine Stier, Simon C. Griffith

## Abstract

Understanding genetic incompatibilities and genetic introgression between incipient species are major goals in evolutionary biology. Mitochondrial genes evolve rapidly and exist in dense gene networks with coevolved nuclear genes, suggesting that mitochondrial respiration may be particularly susceptible to disruption in hybrid organisms. Mitonuclear interactions have been demonstrated to contribute to hybrid disfunction between deeply divergent taxa crossed in the laboratory, but there are few empirical examples of mitonuclear interactions between younger lineages that naturally hybridise. Here we use experimental crosses and high resolution respirometry to provide the first evidence in a bird that inter-lineage mitonuclear interactions impact mitochondrial aerobic metabolism. Specifically, respiration capacity of the two paternal backcrosses (with mismatched mito-nuclear combinations) differ from one another, although they do not differ to the parental groups or maternal backcrosses as we would expect of mitonuclear disruptions. In the wild hybrid zone between these subspecies the mitochondrial cline centre is shifted west of the nuclear cline centre, which is consistent with the direction of our experimental results. Our results therefore demonstrate asymmetric mitonuclear interactions that impact the capacity of cellular mitochondrial respiration and may help to explain the geographic discordance between mitochondrial and nuclear genomes observed in the wild.

## Introduction

Understanding the factors that obstruct gene flow or propel genetic introgression between biological lineages is a central goal of speciation research[1]. One factor that may be particularly susceptible to disruption in hybrids is the interaction between the mitochondrial and nuclear genomes, which underpin mitochondrial aerobic metabolism[2–4]. Functional mitochondria rely on dense gene networks encompassing genes in both the mitochondrial (N= ∼37 genes) and nuclear (N = ∼1,500 genes) genomes (Figure 1A)[5–7], and these genes (particularly mitochondrial genes) can undergo rapid synonymous change between lineages [8–12]. Following hybridization, novel combinations of mtDNA and nuDNA may disrupt mitochondrial function and thus act as a reproductive isolating barrier.

**Figure 1.**
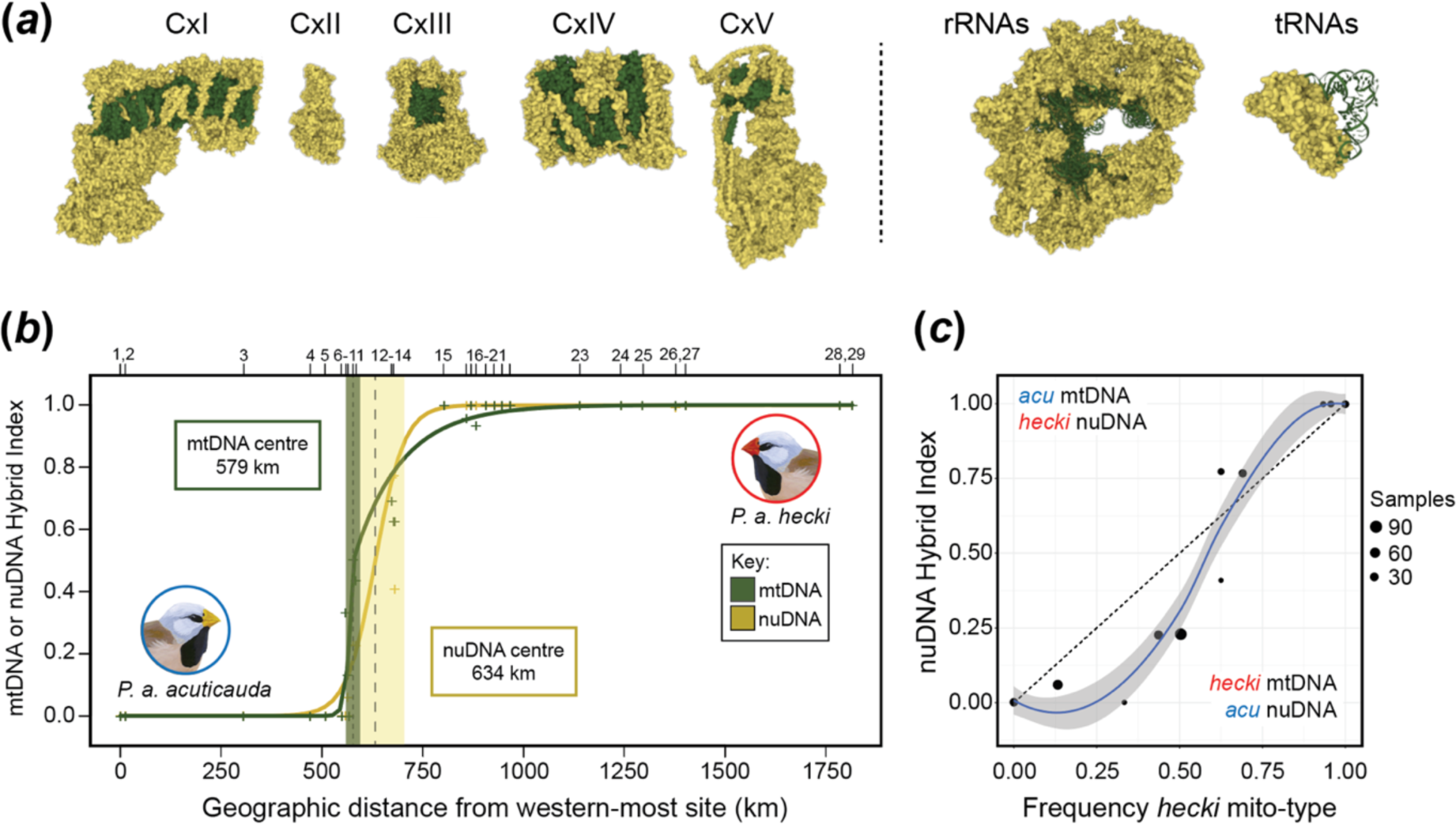
Mitochondrial and nuclear genome interactions in the long-tailed finch. (*a*) Sites of physical interaction within the inner mitochondrial membrane between products encoded by the mitochondrial (green) and nuclear (yellow) genomes. These include proteins that form the electron transport system complexes (CxI-IV) and ATP synthase (CxV), as well as mitochondrial ribosomes and transfer RNA-tRNA sythetases. Complexes are not drawn to scale. The animals and PBD accession numbers for each are CxI (bovine, 5LNK), CxII (chicken, 2H89), CxIII (chicken, 3CWB), CxIV (bovine, 1V45), CxV (bovine, 5ARA – TBC), tRNA (yeast, 4YYE), rRNA (humans, 3J9M). Figure inspired by Hill et al. 2019. **(*b*)** Geographic clines for mtDNA and nuDNA (Z-chromosome) across the range of the long-tailed finch. The mtDNA cline centre is 55 km west of the nuDNA cline centre. The position of 29 sampled populations are shown above the panel and population means for mtDNA and nuDNA are presented as green or yellow crosses, respectively. **(*c*)** The frequency of the *hecki* mito-type against nuDNA hybrid index for 11 populations sampled within the hybrid zone, with a loess curve with 95% CI fitted to the data against a dashed line showing the 1:1 neutral expectation. (***b***) and (***c***) are based on re-evaluated and re-plotted published data from Lopez et al. 2021.

Inter-lineage mitonuclear interactions – i.e., interactions between the mitochondria of one lineage and the nuclear genome of another lineage that occur within the cells of hybrids - have been demonstrated to contribute to hybrid disfunction between deeply divergent taxa crossed in the laboratory [13–17]. There are, however, fewer empirical examples of inter-lineage mitonuclear interactions between younger taxa that naturally hybridise [18–20]. In birds, there is suggestive evidence from hybrid zones that mitonuclear incompatibilities may contribute to reproductive isolation [21–23]. Direct evidence linking hybrid disfunction with mitonuclear incompatibilities is lacking, however, in large part because most birds cannot be readily kept and bred in captivity. This precludes controlled experiments to effectively disassociate the mitochondrial and nuclear genomes. Naturally produced hybrids that have low physiological function may be compromised for various other reasons, such as their parents mating with a heterospecific because they were poor quality individuals [24]. In contrast to genetic incompatibilities of uniform effect on hybrid fitness, interactions of asymmetrical effect [i.e., differential fitness between hybrids based on direction of cross] can result in certain genes introgressing beyond the hybrid zone and geographic discordance between different regions of the genome. In naturally hybridizing systems, geographic discordance between mtDNA and nuDNA is observed at an appreciable frequency but the underlying mechanisms are rarely understood[25]. Discordance is often attributed to demographic disparities or sex-biased asymmetries based mostly on verbal arguments[26], while the functional consequences of mitonuclear interactions such as those described here are rarely considered [25,27,28].

Optimal function of mitochondrial respiration in birds is dependent on 37 genes in the mtDNA, including 13 protein coding genes that encode key components of the well characterised complexes I, III, IV and V in the electron transport system (ETS) on the inner mitochondrial membrane. In addition, avian mtDNA encodes two ribosomal RNAs (rRNAs), 22 transfer RNAs (tRNAs) and a control region (OH). In humans, for example, disfunction of these mitochondrial genes can impact fitness and manifest as mitochondrial diseases and disorders such as neurodegeneration[29,30]. Changes in the amino acid composition of ETS subunits, or genetic substitutions in the tRNA or rRNA machinery that impact the transcription or translation of ETS subunits, can both potentially destabilise the transfer of high-energy electrons causing them to leak from the ETS. Leaking electrons can contribute to the formation of reactive oxygen species (ROS) that can damage cellular macromolecules[30].

The subspecies of the long-tailed finch *Poephila acuticauda acuticauda* and *P. a. hecki* inhabit tropical savannahs across northern Australia and naturally hybridise where their distributions meet on the edge of the Kimberley Plateau in Western Australia[31]. Previous work has estimated they last shared a common ancestor approximately 0.5 million years ago (0.31 – 0.68 Mya, 95% HPD) based on divergence in three mtDNA genes[32]. Mitonuclear discordance has been observed in the wild hybrid zone between these two subspecies, with the centre of the mitochondrial cline being 55 km west of the nuclear cline centre[32], but within the boundaries of the nuclear hybrid zone (Figure 1B & C, [32]). This discordance suggests the eastern mitochondrial lineage (mitotype) has introgressed westward across the hybrid zone. One possible explanation for this apparent introgression is that hybridisation causes novel mitonuclear interactions that asymmetrically reduce the fitness of individuals with a western mitotype and eastern nuclear genome, by adversely impacting aerobic metabolism and therefore ATP synthesis, or through elevated generation of reactive oxygen species (ROS) that results in oxidative stress. The long-tailed finch can be readily kept and bred in captivity, making it an experimentally tractable model system in which to examine the consequences of this potential mitonuclear mismatch.

Here, we test whether inter-lineage mitonuclear interactions impact aerobic metabolism using subspecies of the long-tailed finch. We utilized a cross design that should expose any inter-lineage mitonuclear interactions that exist between the subspecies in the resulting hybrid embryos. We then assessed mitochondrial aerobic respiration capacity using high-resolution respirometry[33].

## Methods

### Study species

Long-tailed finches of each subspecies (*Poephila acuticauda acuticauda* and *P. a. hecki*) used in this study came from captive populations maintained at Macquarie University in Sydney, Australia. Wild caught *P. a. acuticauda* individuals were originally sourced from Mount House (17°02ʹS, 125°35ʹE) and Nelson’s Hole (15°49ʹS, 127°30ʹE) in Western Australia, and wild-caught *P. a. hecki* were sourced from October Creek (16°37ʹS, 134°51ʹE) in the Northern Territory[34,35]. As individuals from these populations show no evidence of genomic admixture between subspecies[31,32] we hereafter refer to them as ‘parental’.

### mtDNA comparison between subspecies

Changes in the mitochondrial genome are required but not sufficient for inter-lineage mitonuclear incompatibility. Even a single substitution in the mtDNA can lead to reproductive isolation (e.g., [48]). To identify substitutions with the potential to contribute to incompatibilities, we sequenced whole mitochondrial genomes from each long-tailed finch subspecies and counted the number of fixed differences and associated amino acid changes within 13 protein coding genes between them. We assembled mitochondrial genomes using whole genome sequence (WGS) data from both long-tailed finch subspecies (*P. a. acuticauda,* N = 14; *P. a. hecki* N = 11) and their sister species the black-throated finch *P. cincta atropygialis* (N = 11). We used bcftools (version 1.9)[49] consensus after calling and incorporating variants relative to the zebra finch reference genome (GCA_003957565.4) in each sample. Ambiguous positions were output with associated IUPAC codes[43]. We exclusively used WGS data derived from DNA extracted from muscle (rather than blood) to ensure that they were enriched for reads from mitochondrial DNA and not reads from the nuclear encoded mtDNA (NUMT) copy. The mitochondrial genome of the zebra finch reference was used as an additional outgroup to help assign substitutions to their lineage of origin. We used the MITOS2 WebServer[50] to extract specific sequences for each mtDNA encoded gene, including all 13 protein coding genes, 22 tRNA and 2 rRNA and the control region. Each gene was aligned using MAFFT (version 7) web-aligner[51] and we manually examined each alignment with Geneious prime (version 2023.1.2) to count and characterize the distribution of fixed differences between each of the three members of *Poephila* (Table S1). For each protein-coding gene, we used the number of fixed differences between subspecies to quantify their absolute divergence (D_XY_).

We next evaluated the extent of non-synonymous change between long-tailed finch subspecies for each protein-coding gene. We generated lineage-specific consensus sequences for each gene and used the vertebrate mitochondrial genetic code to translate these to amino acid sequence. We again used data from the black-throated finch and zebra finch as outgroups to assign amino acid changes to the long-tailed finch subspecies of origin. We classified each amino acid substitution by any associated change in biochemical properties. Finally, we calculated the ratio of the rate of non-synonymous to the rate of synonymous nucleotide substitutions (dN/dS) between long-tailed finch subspecies for each protein-coding gene using the model M0 implemented in the program Easy-CodeML (v1.41)[52].

### Breeding

Embryos were produced by rotating pairs of long-tailed finches through 20 outdoor aviaries (4.1 m long x 1.85 m wide x 2.24 m high), with one pair per aviary containing nest boxes and nesting material. Mealworms, greens, dry seed, and water were provided *ad libitum*. We required females to have been exclusively with the partner for 14 days minimum before any experimental eggs were collected, to avoid embryos being sired by stored sperm from a different male. Nest boxes were checked every day and eggs were collected the day they were laid, given a unique number using a permanent marker, and stored in a ‘soft box’ at cool room temperature (22 °C) for up to seven days before being placed into an incubator (Brinsea Ovation 56 EX, Brinsea Products, Winscombe, U.K.). Eggs were incubated at 37.5°C and 65% humidity for 12 days. On the 12^th^ day (i.e., the day before expected hatch) eggs were removed and measurements made. Working with embryos meant we eliminated the opportunity for the effects of extended parental care by parents of different genetic backgrounds.

### Crossing Design

We produced backcrossed offspring by first breeding F_1_ hybrid females with an unadmixed parental male of each subspecies, sequentially and in random order. Resulting offspring were defined as maternal backcrosses if their father’s lineage *did* match their mother’s mitotype and were defined as paternal backcrosses if father’s lineage *did not* match their mother’s mitotype (Figure 2). Parental pairs of each subspecies were set up to breed at the same time and used as controls.

**Figure 2.**
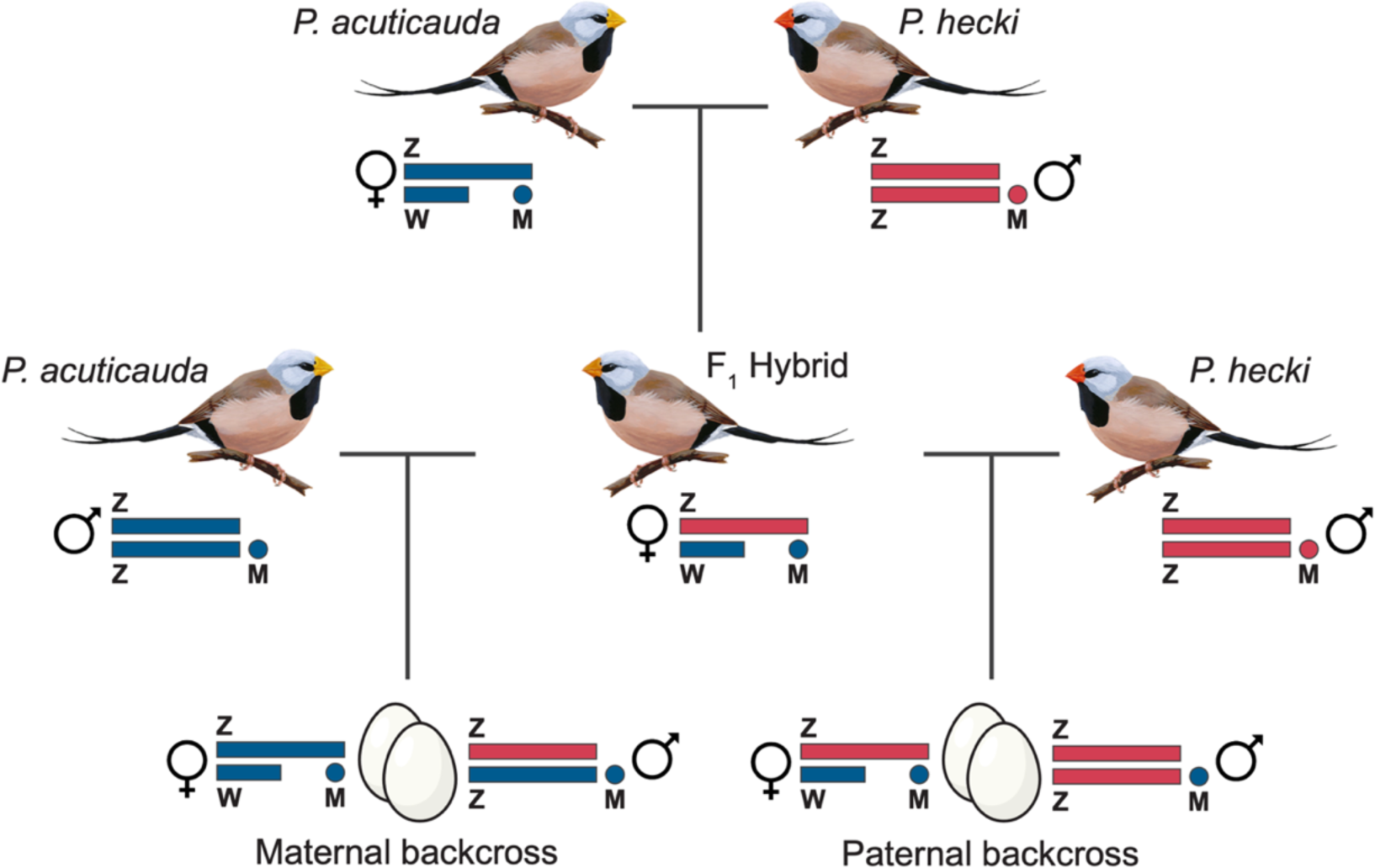
Experimental design used to breed paternal and maternal backcross offspring. The example shown represents the cross between *P. a. acuticauda* females and *P. a. hecki* males used to generate maternal and paternal backcross hybrids carrying *acuticauda* mito-types. The reciprocal cross used to generate *hecki* mito-type carrying backcross hybrids was structured in the same way.

Paternal backcrossing exposes incompatibilities by producing offspring who inherit most of their nuclear genome from a different lineage to that of their mitochondrial genome. This is particularly pronounced in the case of the long-tailed finch, as >98.5% of fixed nuclear differences between subspecies are on the Z-chromosome[36]. In our cross design, paternal backcross offspring inherit Z chromosomes that have had no opportunity for recombination between subspecies. If male, they receive one Z chromosome from their hemizygous F_1_ mother and a Z chromosome of matching subspecies identity from their father. Females have only a single Z chromosome inherited from their father. Paternal backcrosses have mismatched mitochondrial and nuclear genomes, but also have a low level of admixture on autosomes in their nuclear genome, which would otherwise be a potentially confounding source of hybrid disfunction. For this reason, we included maternal backcross individuals as controls, as their level of autosomal admixture is the same as paternal backcrosses, but their mitochondria and Z chromosomes come from the same subspecies (Figure 2). It is worth noting that these several different types of backcrossed hybrids are also highly relevant to the wild hybrid zone, where we previously described that ∼68% (243 of 357) of individuals had some level of admixture[32].

The goal of our cross design was to create embryos in which the mitochondrial haplotype (i.e., mitotype) of the two subspecies was placed in three distinct contexts: 1) a nuclear genetic background exactly matching the mitotype lineage (parental crosses), 2) a nuclear genetic background that mostly did match the mitotype lineage (maternal backcrosses), and 3) a nuclear genetic background that mostly did not match the mitotype lineage (paternal backcrosses). The six resulting groups of embryos were thus 1. Parental *acuticauda* (*acuticauda* mother × *acuticauda* father), 2. Parental *hecki* (*hecki* mother × *hecki* father), 3. Maternal backcross *acuticauda* (F_1_ mother with *acuticauda* mtDNA × *acuticauda* father), and 4. Maternal backcross *hecki* (F_1_ mother with *hecki* mitotype × *hecki* father), 5. Paternal backcross *acuticauda* (F_1_ mother with *acuticauda* mtDNA × *hecki* father), and 6. Paternal backcross *hecki* (F_1_ mother with *hecki* mtDNA × *acuticauda* father). Mothers used in parental crosses only produced a single clutch while F_1_ hybrid mothers used for backcrossing produced two clutches: one with a *hecki* father and one with an *acuticauda* father (however, 4 F_1_ females only produced a clutch with one partner). All fathers were only used for a single clutch.

### Heart rate measurement

After 12 days of incubation each egg was removed from the incubator and immediately placed on a Buddy digital egg monitor (Vetronic Services, Abbotskerswell, Devon, U.K.), to measure embryonic heart rate from within the egg (a proxy for whole body metabolic rate)[37]. A timer was started when the egg was removed from the incubator, and heart rate and corresponding time since removal were recorded at least twice over two minutes. We used these measurements to calculate the rate of cooling and the predicted heart rate halfway through the period (i.e., at 60 seconds). Five individuals where heart rate was only measured once were excluded from this dataset.

### High resolution respirometry strategy

High resolution respirometry was performed with permeabilised tissue samples from these embryos using an Oroboros O2k (Oroboros Instruments, Innsbruck, Austria). High resolution respirometry allowed us to systematically stimulate, uncouple and inhibit certain combinations of mitochondrial complexes (Table 3; Figure S1). We measured the maximum capacity of OXPHOS system when fuelled by electrons either through both complex I, or both I and II in tandem (OXP-I, OXP-I,II), and the maximum capacity of the ETS when accepting electrons from CxII, and both CxI and II in tandem (ETS-I,II, ETS,II). Comparing our experimental groups for these measures reveals whether CxI is compromised. We also assessed the maximum capacity for complex IV in isolation to consume O_2_, a measure not reliant on CxIII, which can reveal if CxIII is compromised. We assessed LEAK-I,II respiration, which reflects respiration not available to fuel ATP synthesis.

These measures enable us to test whether mitonuclear interactions impact mitochondrial aerobic metabolism in long-tailed finch hybrids. More specifically, they allow us to interrogate the source of any mitonuclear disfunction, be it from the two complexes with amino acid differences between subspecies (CxI and III) or from translational issues related to fixed differences in the mtDNA encoded tRNAs or rRNAs. It is possible that some aspect of mitochondrial function other than aerobic respiration is impaired, but that respiration rate is maintained. For this reason we also measured ROS-induced damage. High ROS levels resulting from mitochondrial disfunction can damage macromolecules, cell components and structures[38,39] and ROS-induced DNA damage correlates with shorter lifespan in the closely related zebra finch[40].

### High resolution respirometry technical steps

After the embryonic heart rate was measured, embryos were used for high resolution respirometry. Embryos were euthanised via decapitation and then weighed (Mettler Toledo, PB303-S/FACT, Colombus, Ohio, USA). There was no significant difference between experimental groups in embryonic survival (X^2^_5_ = 4.1, p = 0.53; Figure S3B) or mass at day 12 of incubation (X^2^_5_ = 7.3, p = 0.20; Table 3; Figure S3C). The head and body were then laterally cut in two, and the right-hand side was placed in 400 µl of phosphate buffered saline (PBS), homogenized with three strokes of a teflon7-glass Potter-Elvehjem homogenizer (Wheaton, 5 mL), and stored at -80 for later analyses including DNA extraction for sex determination (described below). The left-hand side was weighed, suspended in a dilution of 100 mg/mL of MiR05 medium (0.5 mM Egtazic Acid (EGTA), 3 mM MgCl2, 50 mM K-lactobionate, 20 mM taurine, 10 mM KH2PO4, 20 mM Hepes, 110 mM sucrose, free fatty acid bovine serum albumin (1 g L-1), pH 7.1;[41,42]); and then homogenized and mechanically permeabilised using a teflon-glass Potter-Elvehjem homogenizer (Wheaton, 5 mL). All work was done on ice. This homogenate was centrifuged for 1 min at 100 g to pellet big tissue, and then 500ul of the supernatant was added to a chamber of an Oroboros O_2_k high resolution respirometer set at 37°C. The respirometer had two chambers and we always ran two samples in parallel. After two samples were loaded and the system had equilibrated (oxygen consumption was stable), 2 µl of digitonin (10 mg/mL) was added to ensure cells were fully permeabilised. We injected 5 µl pyruvate (5mM final), 5 µl malate (2mM final), 10 µl glutamate (10mM final) as electron donors for CxI, and then measured oxidative phosphorylation when only CxI was receiving electrons (OXPHOS-I) by adding 5 µl adenosine diphosphate (ADP, 1.25 mM final). We next measured coupled respiration capacity through both CxI and II (OXPHOS-I,II) by adding 25 µl succinate (10 mM final). We then determined O_2_ consumption associated with proton leak when electrons were being received by both CxI and II (LEAK-I,II) by adding 5 µl oligomycin (10 uM final). We next estimated the maximum capacity of the electron transport system (ETS-I,II) by titrating the mitochondrial uncoupler carbonyl cyanide m-chlorophenyl hydrazine (CCCP; at 2 mM concentration) by first one injection of 3 µl, followed by 1 µl steps. We next estimated the ETS capacity when only receiving electrons through CxII (ETS-II) by adding 1 µl rotenone (0.5 uM final). We then measured non-mitochondrial O2 consumption by injecting 1 µl of antimycin A (2.5 uM final) to inhibit CxIII. Finally, we estimated the respiration capacity of CxIV directly by first adding 5 µl TMPD (0.5 mM final) and 5 µl ascorbate (2 mM final), and then calculating the background chemical respiration rate by adding 50 µl azide (100 mM final). Across all runs the chambers were opened consistently at the same stages for reoxygenation. After each run 1 ml of the solution was taken from each respirometer chamber and frozen at -80 °C, and at the conclusion of all measures we performed a Pierce BCA protein quantification assay on this stored solution (ThermoFisher, Scientific, Waltham, MA, USA). Protein quantity was included as a covariate in statistical models (below) to account for slight differences in the amount of biological material in each sample. In most cases a different embryo sample was run in each chamber, but in 40 cases the same embryo was run in duplicate to assess how repeatable these measurements were (Table 3). When two different samples were run, the first was sitting in the respirometer for some time before the protocol was begun, and to account for any variability introduced by this we included a covariate in our models that was the length of time between when the embryo sample entered the chamber and when the measure was made.

### Molecular sexing

We extracted DNA from the homogenized right-side of all embryos using a Gentra PureGene kit (Qiagen, Valencia, CA, USA) following manufacturer instructions. We sexed embryos by amplifying an intronic portion of the CHD gene that differs in size between the Z-linked and W-linked copy following published protocols[32,43]. Gel electrophoresis of the PCR product revealed the sex of the embryo as either male (one band, homozygous, ZZ) or female (two bands, heterozygous, ZW).

### mtDNA copy number

We estimated mitochondrial density based upon mtDNA copy number for all samples using real-time quantitative PCR (qPCR) analysis following protocols established for passerine birds (Stier et al. 2019, 2022). While the amount of mtDNA in a mitochondrion can vary, mtDNA copy number is a reliable proxy of mitochondrial density. We estimated the relative mtDNA copy number by measuring the amount of mtDNA relative to the nuDNA for each sample by qPCR on a CFX96 Real-Time System (BIO-RAD). We used RAG1 as a representative single-copy nuclear gene (F: GCAGATGAACTGGAGGCTATAA, R: CAGCTGAGAAACGTGTTGATTC) and used COX1 as our representative mitochondrial gene (F: TCCTAGCCAACTCCTCAC, R: CCTGCTAGGATTGCAAAT). A complete nuclear copy of the mitochondrial genome (i.e., a NUMT) exists in Estrildid finches; for example, in the zebra finch reference genome (GCA_003957565.4) at chr2:72232845-72249691. To minimize the risk of NUMT interference in estimating mtDNA copy number, we designed our COX1 primers to perfectly match the mitochondrial sequence observed in *Poephila*. Moreover, our use of embryo tissue is less likely to result in amplification of the NUMT as gDNA from muscle tissue has a much lower nuclear:mitochondria ratio than gDNA from blood[44]. We further verified that COX1 amplification resulted in a single product of the expected size on an agarose gel.

Following Stier et al. (2022), qPCR reactions had a total volume of 12 µl that included 6 ng of DNA, had PCR primers with concentrations of 200 nM and 6 µl of Absolute Blue qPCR Mix SYBR Green low ROX (Thermos Scientific). The qPCR conditions were: 15 mins at 95°C, 40 cycles of 15 s at 95°C, 30 s at 58°C, 30 s at 72°C. Samples were run in triplicate for each gene (RAG1 and COX1) across 5 plates, ten samples were pooled and included on every plate as a reference, and 20 samples were included on two separate plates. Amplicon efficiency was estimated from a standard curve of the reference between 1.5 to 24 ng (1.5, 3, 6, 12, 16, 24) run on one plate. The mean reaction efficiencies were 92.1% for RAG1 and 82.3% for COX1. To calculate relative mtDNA copy number for each sample we used the following formula: (1+Ef_COX1_)^ΔCqCOX1^/(1+Ef_RAG1_)^ΔCqRAG1^, with Ef as the amplicon efficiency, ΔCq as the difference in Cq between the reference and focal samples. Some samples were run twice on separate plates, with an inter-plate repeatability of 0.74 (95% C.I. [0.49, 0.88]), and based on triplicates intra-plate repeatability was 0.94 (95% C.I. [0.93, 0.95]). Seven samples were considered as outliers (Cq RAG1 < Cq COI) and removed from analysis.

### Oxidative Damage assay

8-OHdG (8-hydroxy-2’-deoxyguanosine) is a predominant form of ROS-induced oxidative lesion that has been widely used as a biomarker for oxidative damage[45], including alongside measures of respiration capacity in birds[46,47]. Here we quantified levels of 8-OHdG in the DNA extracted from embryos using a competitive immunoassay (300 ng DNA, EpiQuick 8-OHdG DNA Damage Quantification Direct Kit Colorimetric, Epigentek, USA) following the manufacturer recommendations. This was run on a subset of samples (N = 182) chosen blind to respiration capacity to have roughly balanced numbers from each experimental group (N = 28-33 samples per group). The intra-plate coefficient of variation based on duplicates was 6.3 ± 0.83. The inter-plate coefficient of variation based on four samples repeated over all three plates was 22.7 ± 4.05, so we included plate ID as random effect in our analyses.

### Statistical analyses

For statistical analyses we used R version 4.1.2 (R Core Team 2020) using RStudio version 1.2.5033 (RStudio Team, 2020) for the graphical interface. We used the lme4 and lmerTest packages[53,54] to run linear mixed models to test whether our measures of respiration capacity differed between the six experimental groups (parental *acuticauda*, parental *hecki*, maternal and paternal backcross with acuticauda mtDNA, maternal and paternal backcrosses with hecki mtDNA). These measures were each included as the dependent variable in separate linear mixed models, in which the fixed effects were experimental group, embryo sex, protein quantity in the chamber, and the length of time after embryo entered the oroboros chamber when the measure was made. In each we tested for an interaction between experimental group and embryo sex, and if this was not significant the interaction term was removed (but left in the model if significant). In each model parental pair ID was included as a random effect, to account for multiple members of the same clutch being included. Where there was a significant effect of experimental group, we ran Tukey post-hoc comparisons using the emmeans package[55] to assess specifically which groups differed from one another. Model assumptions were checked using the DHARMa package in R[56].

We similarly ran linear mixed models to test whether heart rate, oxidative damage and mitochondrial copy number differed between our experimental groups, but in these cases the fixed effects were experimental group, embryo sex, and for the heart rate model embryo mass, and the random effect was still parental pair ID. Oxidative damage and mitochondrial copy number were log-transformed to better meet model assumptions. An interaction between experimental group and sex was also included for each model, but then removed if not significant. In the mitochondrial copy number and oxidative damage models, plate ID were additionally included as random effects. Embryo ID was also initially included as a random effect in the mitochondrial copy number and oxidative damage models, but it explained close to zero variance so was removed for the final models, likely because very few individuals were measured twice (10 for mitochondrial copy number, 21 for oxidative damage). A small number of values for mitochondrial copy number were extremely high (N = 10, total N = 211), well above the normally distributed other samples and were likely technical errors, and so were excluded to facilitate modelling. When plotted they were distributed roughly evenly between experimental groups so their inclusion would be unlikely to change the results anyway.

## Results

We found a low level of divergence in mtDNA between subspecies, with fixed differences observed in only 0.9% of positions comprising the protein-coding genes of the electron transport system (107 of 11,361 bp total; Table 1). Absolute divergence (D_XY_) was highly variable, however, across protein-coding genes, ranging from 0% (ATP8) to 3.3% (NAD6), and between complexes: divergence was 2.1-fold higher on average for genes in complex I than in complex IV (Table 1). We observed a total of seven fixed amino acid differences between subspecies, five differences in four genes in complex I, and two amino acid differences in the cytochrome b (COB) gene in complex III. There were no amino acid differences between the subspecies in either complex IV or complex V. The inferred mitochondrial genome-wide dN/dS ratio between long-tailed finch subspecies is well below 1 at 0.0168, suggesting that purifying selection is the dominant force shaping mitochondrial evolution in *Poephila* (Table 1). Of the five genes with observed non-synonymous differences between subspecies, dN/dS ranged from 0.0127 (*NAD5*) to 0.1142 (*NAD3*).

**Table 1.** Mitochondrial protein-coding divergence between long-tailed finch subspecies. . The 13 protein-coding genes comprising the electron transport system (ETS) are organized by complex. Fixed differences between subspecies were inferred based on comparison of full mitochondrial genomes of *Poephila a. acuticauda* (N = 14) and *P. a. hecki* (N = 11) with D_XY_ calculated as the absolute divergence between members of each subspecies. Estimates of dN/dS were calculated for each gene using the model M0 implemented in the program Easy-CodeML (v1.41; Gao et al 2019). The combined estimate of dN/dS was inferred based on net dN (0.0114) and dS (0.6802) across all protein-coding genes.

| Complex | Gene | Fixed differences | $D_{xy}$ | dN/dS |
| --- | --- | --- | --- | --- |
| Complex I | *NAD1* | 12 | 0.012 | 0.0001 |
|  | *NAD2* | 8 | 0.008 | 0.0001 |
|  | *NAD3* | 3 | 0.009 | 0.1142 |
|  | *NAD4* | 9 | 0.007 | 0.0599 |
|  | *NAD4L* | 5 | 0.017 | 0.0001 |
|  | *NAD5* | 17 | 0.009 | 0.0127 |
|  | *NAD6* | 17 | 0.033 | 0.0210 |
| Complex III | *COB* | 10 | 0.009 | 0.0574 |
| Complex IV | *COX1* | 9 | 0.006 | 0.0001 |
|  | *COX2* | 4 | 0.006 | 0.0001 |
|  | *COX3* | 3 | 0.004 | 0.0001 |
| Complex V | *ATP6* | 10 | 0.015 | 0.0001 |
|  | *ATP8* | 0 | 0.000 | 0.0001 |
| **Combined** |  | **107** | **0.009** | **0.0168** |

In the non-coding region, we observed a further six fixed differences between subspecies in the small and large rRNA genes (four and two differences, respectively) and a total of four fixed differences in four mitochondrial tRNAs (Table S1). These non-coding differences provide additional potential sites of mitonuclear incompatibilities that could impact mitochondrial function via the transcription, translation, or replication of mtDNA [e.g., 41]. On the other hand, any protein-protein mitonuclear incompatibilities in this system should be restricted to impacting the function of complexes I or III.

Polarization inference from the use of outgroup taxa revealed that of the seven fixed amino acid differences between subspecies, six have occurred in the *hecki* lineage (Figure 3; Table 2). Three of the amino acid substitutions derived within *hecki*, and the single amino acid change derived in *acuticauda*, result in a change from a polar to a non-polar amino acid (Table 2). In the non-protein coding genes (rRNAs, tRNAs, OH) there was a roughly even spread of derived DNA sequence changes between lineages (6 *acuticauda* and 4 *hecki,* Table S1 and S3).

**Figure 3.**
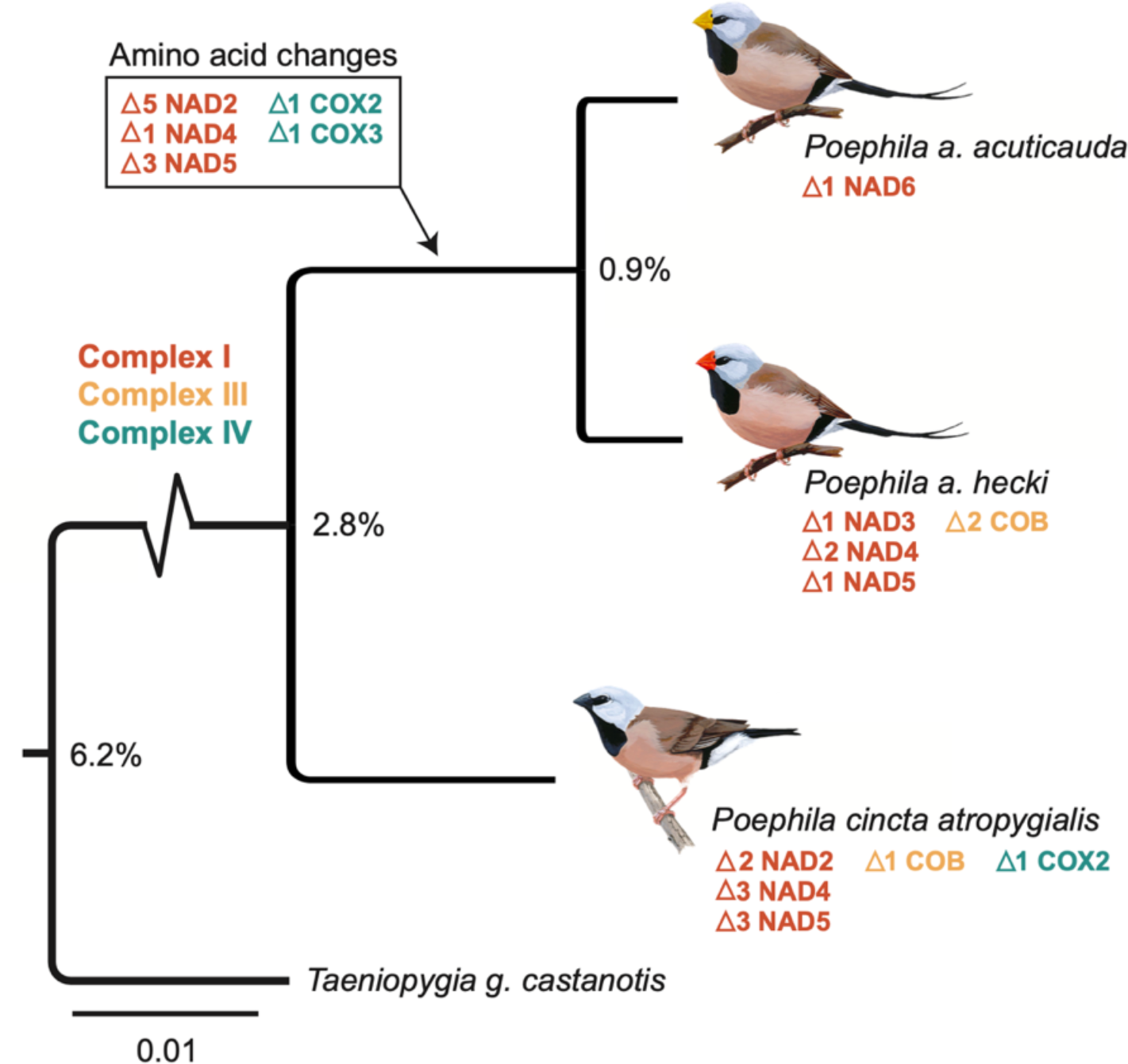
Evolutionary history of mtDNA protein-coding divergence between long-tailed finch subspecies. A maximum likelihood tree was generated based on an aligned concatenation of consensus sequences for all 13 protein-coding genes using W-IQ-TREE [1] under the best-fit substitution model: TPM2u+F+I. The zebra finch (NCBI accession: DQ422742) and black-throated finch samples were used to assign all nucleotide substitutions and any associated amino acid changes to their branch of origin. Branch lengths are proportional to the number of substitutions. Divergence estimates at each node represent the total number of fixed differences between species in protein-coding genes divided by the total number of protein-coding bases (Table S2). The number of amino acid changes identified within each gene are color-coded by ETS complex: red for complex I, yellow for complex III, and blue-green for complex IV.

**Table 2.** mtDNA amino acid differences between long-tailed finch subspecies. . Fixed differences were assigned to their lineage of origin using mitochondrial genomes of the black-throated finch (*Poephila cincta atropygialis*; N = 11) and the zebra finch (*Taeniopygia guttata*; N = 1) as outgroups.

| Complex | Gene | Mutation | Biochemical Change | Origin |
| --- | --- | --- | --- | --- |
| Complex I | *NAD3* | T9A | polarity change (+ > -) | *hecki* |
|  | *NAD4* | T96A | polarity change (+ > -) | *hecki* |
|  | *NAD4* | T398A | polarity change (+ > -) | *hecki* |
|  | *NAD5* | A16V | none | *hecki* |
|  | *NAD6* | S127L | polarity change (+ > -) | *acuticauda* |
| Complex III | *COB* | S358T | none | *hecki* |
|  | *COB* | I372M | none | *hecki* |

We observed a significant difference between the two paternal backcross groups for all five measures of aerobic respiration capacity (except borderline non-significant for females for OXP-I,II, p = 0.064), whereby individuals with *acuticauda* mtDNA and *hecki* nuDNA had lower respiration capacity than did individuals with *hecki* mtDNA and *acuticauda* nuDNA (Table 3; Figure 4, Figure S2). There were no other differences between experimental groups that were as consistent across the five measures of aerobic capacity we measured (Table 3). For the measure ETS-I,II the paternal backcross *hecki* mitotype was significantly higher than the maternal backcross *acuticauda* mitotype, and for OXP-I females the parental group *hecki* was significantly higher than the paternal backcross *acuticauda* mitotype group (Table 3).

**Table 3.** Model output comparing the measures of mitochondrial function and physiology measures among the six experimental groups.

| Respiration measure | ETS Cxs included | Description of measure | Repeatability (SE) from 40 samples run in duplicate | Cross*Sex | Sex (if split model due to interaction) | Cross | Sex | Protein Qty | Length of time | Var. | MargR<sup>2</sup> | ICC Pair ID | CondR<sup>2</sup> | G5<G6 | G6>G3 | G2<G6 | G2>G5 | G5<G4 |
| --- | --- | --- | --- | --- | --- | --- | --- | --- | --- | --- | --- | --- | --- | --- | --- | --- | --- | --- |
|  |  |  |  | X2 (DF) p-value |  | X2 (DF) p-value | X2 (DF) p-value | X2 (DF) p-value | X2 (DF) p-value |  |  |  |  | Showing p-values for p < 0.1. |  |  |  |  |
| OXP-I,II | I,II,III,IV,V | Coupled respiration | 0.94 (0.02) | 11.5 (5) 0.043 | Female | 12.73 (5) 0.026 | NA | 2.59 (1) 0.11 | 8.68 (1) 0.003 | 197 | 0.62 | 0.37 | 0.99 | 0.064 |  |  | 0.072 |  |
|  |  |  |  |  | Male | 13.87 (5) 0.016 | NA | 7.98 (1) 0.005 | 9.31 (1) 0.002 | 240 | 0.44 | 0.55 | 0.99 | 0.042 |  | 0.093 |  |  |
| ETS-I,II | I,II,III,IV | Uncoupled respiration | 0.90 (0.03) | 9.78 (5) 0.08 |  | 20.15 (5) 0.001 | 2.3 (1) 0.13 | 4.06 (1) 0.044 | 26.1 (1) <0.001 | 289 | 0.53 | 0.46 | 0.99 | <0.001 | 0.034 |  |  |  |
| OXP-I | I,III,IV,V | Coupled respiration | 0.69 (0.09) | 11.97 (5) 0.035 | Female | 15.4 (5) 0.009 | NA | 3.29 (1) 0.07 | 25.2 (1) <0.001 | 206 | 0.79 | 0.20 | 0.99 | 0.029 |  |  | 0.036 |  |
|  |  |  |  |  | Male | 14.9 (5) 0.010 | NA | 6.67 (1) 0.010 | 33.11 (1) <0.001 | 224 | 0.63 | 0.36 | 0.99 | 0.030 |  | 0.052 |  |  |
| ETS-II | II,III,IV | Maximum capacity of the ETS | 0.96 (0.01) | 7.1 (5) 0.2 |  | 19.6 (5) 0.002 | 1.53 (1) 0.22 | 5.04 (1) 0.02 | 1.17 (1) 0.28 | 55 | 0.39 | 0.58 | 0.98 | 0.001 | 0.065 |  |  |  |
| CxIV | IV | Maximum respiration | 0.93 (0.02) | 8.7 (5) 0.12 |  | 19.8 (5) 0.001 | 1.5 (1) 0.22 | 7.8 (1) 0.005 | 2.14 (1) 0.14 | 302 | 0.41 | 0.58 | 0.99 | 0.001 |  |  |  | 0.056 |
| LEAK-I,II | I,II,III,IV | Respiration linked to proton leak | 0.89 (0.03) | 9.6 (5) 0.09 |  | 17.3 (5) 0.004 | 2.4 (1) 0.12 | 8.6 (1) 0.003 | 0.3 (1) 0.58 | 1.4 | 0.19 | 0.36 | 0.55 | 0.003 |  |  |  |  |
| Measure |  |  |  | Cross*Sex | Sex (if split model due to interaction) | Cross | Sex | Embryo Mass |  | Var. | MargR<sup>2</sup> | ICC Pair ID | CondR<sup>2</sup> |  |  |  |  |  |
| Embryo Mass |  |  |  | 8.6 (5) 0.13 |  | 7.3 (5) 0.20 | 0.01 (1) 0.92 | NA |  | <0.001 | <0.001 | <0.001 | <0.001 |  |  |  |  |  |
| Embryo Survival |  |  |  | NA |  | 4.1 (5) 0.53 | NA | NA |  | 0.09 | 0.02 | 0.24 | 0.27 |  |  |  |  |  |
| Heart rate (proxy for whole body metabolism) |  |  |  | 15.2 (5) 0.01 | Male | 7.8 (5) 0.17 | NA | 23.5 (1) <0.01 |  | 156 | 0.41 | 0.59 | 0.99 |  |  |  |  |  |
|  |  |  |  |  | Female | 6.3 (5) 0.28 | NA | 8.3 (1) 0.004 |  | 78 | 0.20 | 0.80 | 0.99 |  |  |  |  |  |
| Oxidative DNA oxidative damage (log transformed) |  |  |  | 8.3 (5) 0.14 |  | 8.6 (5) 0.12 | 3.0 (1) 0.08 | NA | ICC Plate: 0.008 | 0.019 | 0.006 | 0.012 | 0.026 |  |  |  |  |  |
| mtDNA copy number (log transformed) |  |  |  | 2.2 (5) 0.82 |  | 20.9 (5) <0.001 | 0.17 (1) 0.69 | NA | ICC Plate: 0.00 | 0.03 | 0.01 | 0.02 | 0.03 | G5<G6 0.005 | G1<G6 0.093 | G3<G6 0.090 |  |  |

**Figure 4.**
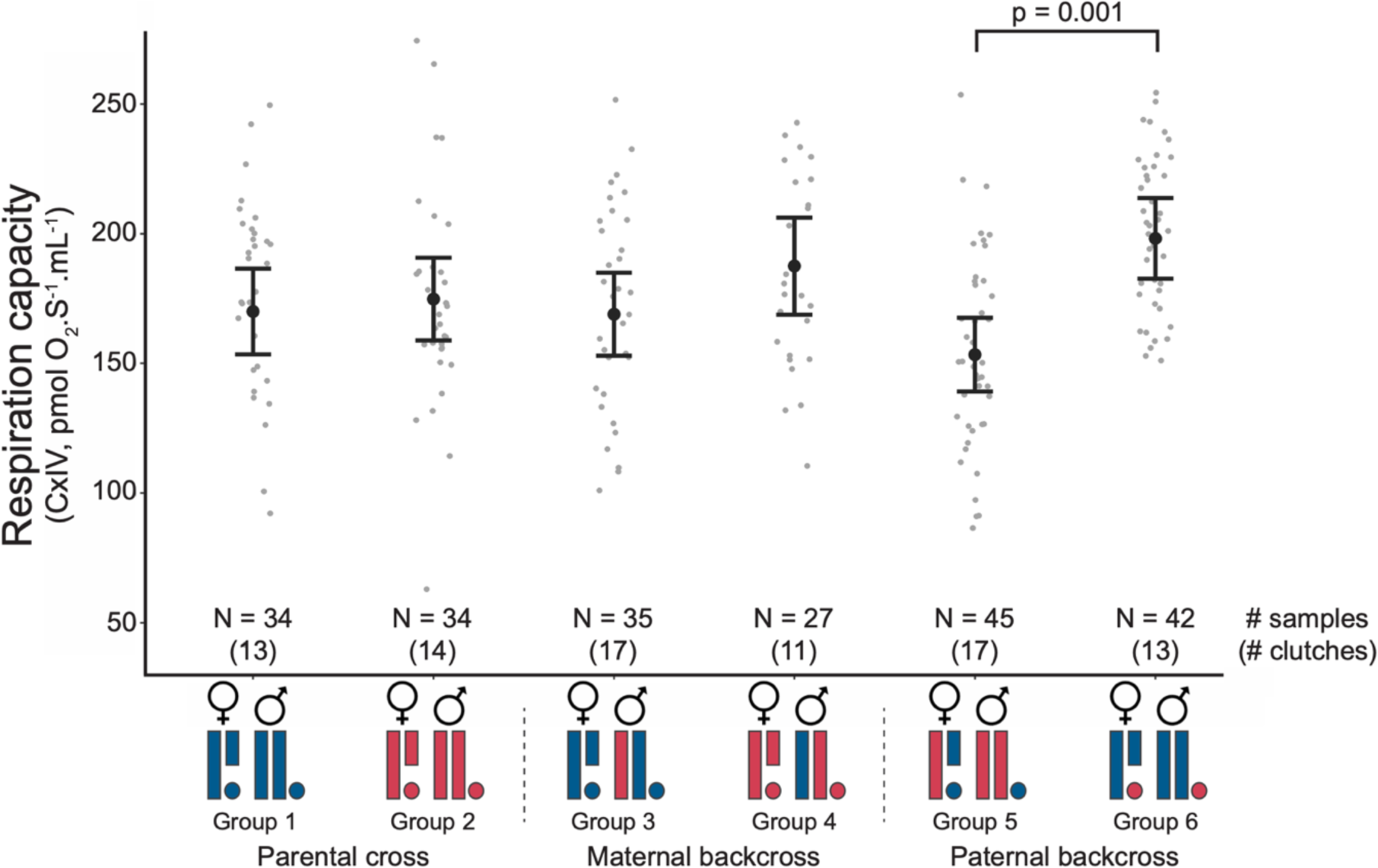
Mitochondrial and nuclear genomes between the long-tailed finch subspecies interact to impact mitochondrial respiration capacity. The respiration capacity of our experimental groups, with grey datapoints indicating raw data (corrected for amount of protein), and black mean and 95% CIs reflect model output (accounting for clutch ID, protein quantity, length of time in chamber, sex). Most measures of mitochondrial respiration capacity had this same pattern of differences among the groups (Table 3; Figure S2).

We did not find a difference between groups in oxidative damage (Table 3, Figure S3D). We also found no evidence that this difference in aerobic respiration capacity between paternal backcross groups was driven by different quantities of mitochondria per cell, with the relative mtDNA copy number not significantly differing between the two paternal backcross groups, or between them and other groups (Table 3) and so likely not driving the aerobic respiration capacity results. We also found no difference between experimental groups in heart rate on day 12 of incubation immediately after being removed from the incubator and so no evidence that whole body metabolism is impacted by hybrid cross (Table 2, Figure S3C).

## Discussion

We found evidence that an interaction between the mitochondrial and nuclear genomes of long-tailed finch subspecies impacted aerobic respiration capacity in hybrids, despite the low level of divergence in mtDNA between them (<1%). These observations identifying the effect of mitonuclear incompatibilities on aerobic respiration have important implications for both mtDNA introgression and reproductive isolation in the long-tailed finch system. Positive selection can drive allelic introgression across a hybrid zone, as can neutral processes (such as range expansion). In the long-tailed finch hybrid zone there are high levels of admixture (∼68% of individuals have some degree of mitonuclear mismatch), and so selection will be acting on each mitotype in several different nuclear contexts, akin to those generated through our experimental approach. Our primary result suggests that there is no difference in mitochondrial aerobic metabolism between the mitotypes in the pure parental context or in the maternal backcross individuals, but that the *acuticauda* mitotype has reduced respiration capacity in the paternal backcross context compared to the *hecki* mitotype in the paternal backcross context. In the regime of widespread admixture within the hybrid zone, negative selection on the *acuticauda* mitotype might result in the *hecki* mitotype increasing in frequency and introgressing westwards. This is indeed what is observed in the wild long-tailed finch hybrid zone, where the centre of the mitochondrial cline is displaced 55 km west of the nuclear cline centre (Figure 1B & C). Respiration capacity reflects the upper limit of mitochondrial aerobic respiration. Therefore, the reduced aerobic capacity observed for *acuticauda* mtDNA paternal backcross individuals is expected to be detrimental, particularly for energetically demanding tissues and life-stages. This suggests that the physiological effects observed in this experiment may be a good proxy for the selection that is occurring in the wild.

We did not, however, find strong evidence that mitonuclear incompatibilities reduce hybrid fitness compared to parental individuals. There were few differences between our paternal backcross groups and either pure parental individuals or maternal backcross groups (Table 3; Figure 4; Figure S2). A conservative interpretation would suggest that incompatibilities between the mitochondrial and nuclear genomes of these subspecies is not a key component of reproductive isolation between them, which might instead be linked to other incompatibilities such as those involving chromosomal inversions [31]. Our lack of difference between parental and paternal backcross groups are in stark contrast to results from comparable experiments using deeply diverged lineages crossed in the lab (e.g., with mitochondrial divergence >10%), where paternal backcrossed individuals had dramatically impacted mitochondrial function, survival, or fecundity [13,14,62,63]. There are however some lines of evidence suggestive of nascent mitonuclear incompatibility in this study. For a few respiration measures, there were significant – or borderline significant – differences between paternal backcross and maternal backcross groups; for ETSI,II (Figure 4) and ETS-II *hecki* paternal backcrosses had higher respiration capacity than did *acuticauda* maternal backcrosses, and for CxIV the *acuticauda* paternal backcrosses had borderline significantly lower respiration capacity than *hecki* maternal backcrosses (Table 3; Figure S2). Furthermore, while not statistically significant, the mean respiration values of both paternal backcross groups were shifted away from the parental groups, which may reflect divergence from the species-specific optima. It is possible that we may simply not have had enough statistical power to detect weak effects. Mitonuclear disfunction may only become apparent in specific tissue types with high baseline energy consumption (e.g., cardiac or nervous system tissue) or under stressful conditions with heightened energetic demands (e.g., during reproduction or thermal stress). Thus, our observations do not preclude a role for mitonuclear incompatibilities as a reproductive isolating barrier at this early stage of population divergence.

We were interested in distinguishing what mitonuclear components were likely interacting to impact mitochondrial aerobic respiration. We wanted to know whether the observed differences in aerobic respiration between paternal backcross groups was the result of specific protein-protein interactions in complexes I and III or if instead they were due to interactions involving mtDNA encoded tRNA or rRNA. The difference between the two paternal backcross groups was consistent across all measures, which included the CxIV respiration measure that is not dependent on other complexes and there are no fixed differences in CxIV amino acids between subspecies. This suggests that the causal mitonuclear interaction may not be between proteins encoded by the two genomes but may instead be between either mtDNA encoded tRNA and nuDNA encoded aminoacyl tRNA synthetases, or between mtDNA encoded rRNA and nuDNA encoded ribosomal proteins. These interactions would impact the translation of mtDNA and therefore the production of all mtDNA encoded ETS complexes (CxI, III, IV, and V).

There was also a significant sex difference in the *hecki* subspecies for two measures (OXP-I,II and OXP-I), whereby females had higher respiration capacity than males (Table 3; Figure S2). As a result, female parental *hecki* individuals had higher respiration capacity than *acuticauda* mtDNA parental backcross individuals, and male parental *hecki* individuals had lower respiration capacity than did *hecki* mtDNA parental backcross individuals. It is unclear to us why there is a sex difference in respiration capacity for *hecki* individuals but not *acuticauda* or backcrossed individuals. This may just be a false positive, as we have run many statistical tests in this study, but it may be biological; sex differences in mitochondrial function have been observed in the literature, although there is no apparent consistent difference[64]. Furthermore, as embryos we would not expect dramatic sex differences in metabolism, except perhaps the gonads. ‘Mother’s curse’ is a phenomenon that can result in a sex effect due to mitonuclear interactions[65–67], but it predicts that detrimental impacts of mtDNA on males and resulting sex differences are exposed with heterospecific nuDNA but not homospecific nuDNA (due to nuDNA coevolution to rescue function), whereas in the present study we found the opposite; that seemingly detrimental impacts of mtDNA only occurs in homo(sub)specific nuDNA context.

We did not find evidence that the observed effects on cellular aerobic respiration capacity between paternal backcross groups affected our proxies for whole-organism metabolic rate or the level of oxidative damage. The inter-lineage mitonuclear interactions may therefore be impacting fitness through other pathways or under specific circumstances, for example in energetically demanding tissues and life-stages. Given the experimental context it would also be valuable to confirm the whole-body result with more direct measures of metabolism (e.g., open-flow respirometry[68]). This is because other differences between experimental groups may undermine how well heart rate reflects differences in whole-body metabolism, for example at low incubation temperature, Japanese quail heart rate decreases[69] but heart volume increases to compensate (A. Stier, unpublished data).

Purifying selection appears to be the dominant force in mtDNA evolution in *Poephila* (the mitochondrial genome-wide dN/dS ratio is well below 1 at 0.0168). The observed amino acid differences between long-tailed finch subspecies are thus notable, with all six *hecki*-derived amino acid changes are in genes within complexes I (four AAs) and III (two AAs) suggesting some degree of inter-molecular epistasis might exist between these substitutions. Moreover, in one of the genes with multiple non-synonymous substitutions (COB), amino acid changes were located <15 codon positions apart, a possible signature of intra-molecular epistasis [57,58]. Taken together this may constitute evidence of adaptive evolution within *hecki*, as mitochondria can contribute to local adaptation (e.g., to differences in climate)[59–61], but to confirm this requires further investigation.

In conclusion, we here demonstrate that functional consequences of interactions between mtDNA and nuDNA are possible even at a low level of divergence (<1% mtDNA), and the resulting impact on fitness between the two groups with mitonuclear mismatch in this system may be strong enough to result in asymmetrical mitochondrial introgression across a hybrid zone. Such mitonuclear interactions may be widespread, as most avian lineages with stable hybrid zones have equal or greater divergence. Researchers considering cases of mitonuclear discordance (or cyto-nuclear discordance more generally) should consider that functional consequences of mitonuclear interactions may be at play[23,70]. Identifying the specific genes involved in mitonuclear interactions in the long-tailed finch would be a valuable direction for future research, as this will facilitate testing the molecular pathways involved in mitonuclear interactions and the evolutionary processes that led to them (e.g., rapid evolution)[20,63,71–74]. This sensitivity of mitochondrial function to genetic variation, their centrality to organismal function (generating 90% of cellular energy in animals[75]), and the broad range of taxa that rely on mitochondria for energy production support suggestions that mitonuclear interactions may be a powerful force acting on hybridising animals[76–79].

## Ethics statement

This experiment was approved by the Macquarie University Animal Ethics Committee (reference number: 2020/021) and all experiments conform to the relevant regulatory standards.

## Data Accessibility and Benefit Sharing

Data from the respiration capacity, oxidative damage, heart rate and mitochondrial copy number, the R scripts for statistical analyses are available on the Open Science Framework (https://osf.io/f652p/?view_only=30142041d5d74dd7bd904ddac0186a41). Mitochondrial sequences for this project will be deposited in GenBank’s nucleotide archive at time of publication. The WGS data used to assemble mitogenomes is part of another study currently in prep and will be made available when that work is published.

## Acknowledgements

The authors would like to thank Riccardo Ton and Hector Pacheco Fuentes for providing assistance setting up the Oroboros O_2_k, and Macquarie Animal Research Services (M.A.R.S.) staff for providing daily bird husbandry.

## Author contributions

Conceptualization, C.S.M., S.C.G.; Methodology, C.S.M., D.M.H., A.S., S.C.G; Investigation, C.S.M.; Data analysis and interpretation, C.S.M., D.M.H., A.S.; Resources, S.C.G.; Writing – Original Draft, C.S.M.; Writing – Review & Editing, C.S.M, D.M.H., A.S., S.C.G.; Visualization, C.S.M., D.M.H.; Supervision, S.C.G.; Funding Acquisition, C.S.M., D.M.H., S.C.G.

## Supplementary Figures

**Figure S1.**
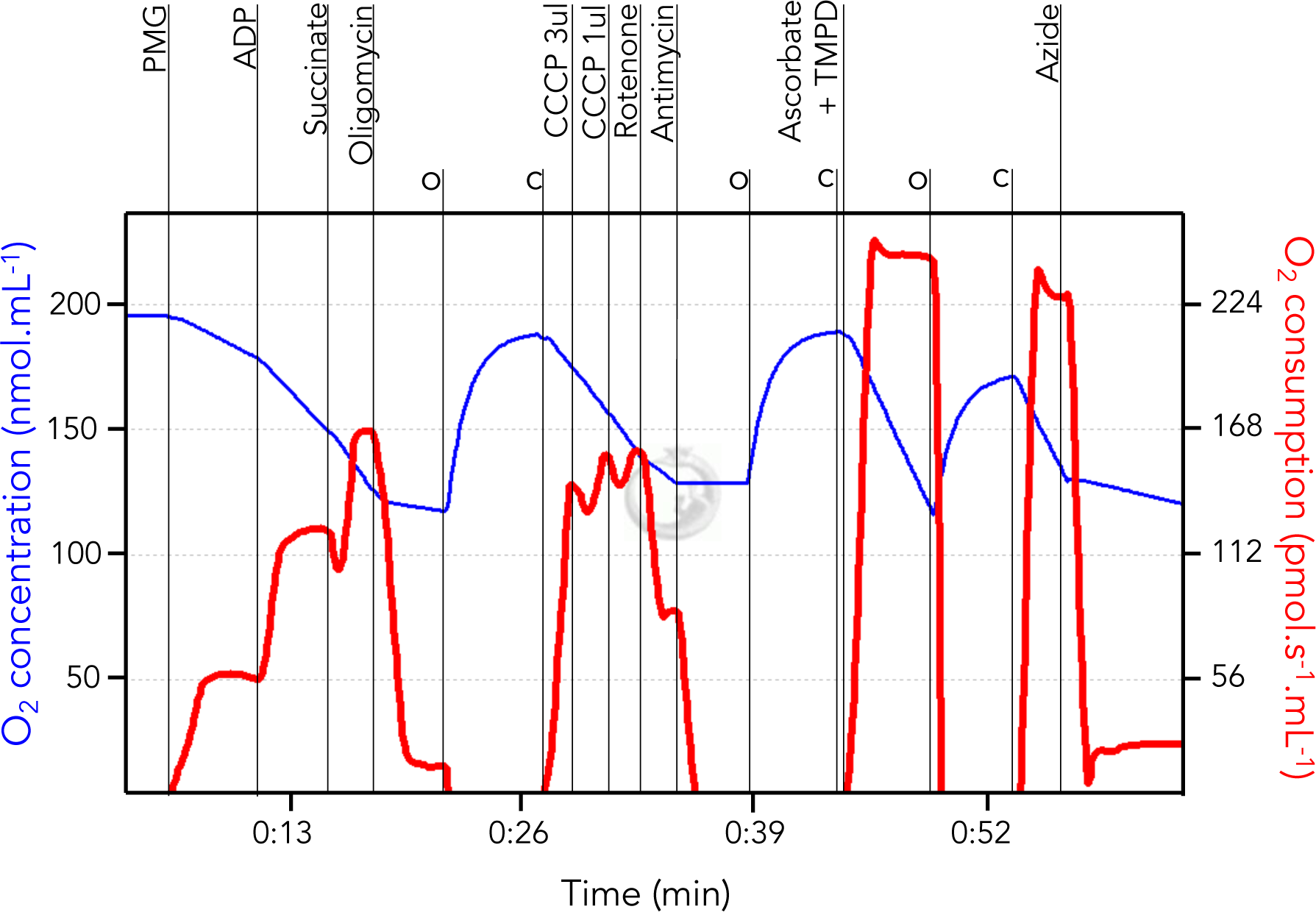
Example of reading from measurement of mitochondrial respiration capacity. Oxygen concentration (blue line) and oxygen consumption (red line) following the injection of substrate, uncoupler, inhibitor titrations (SUIT) injections (long vertical lines). Chambers are opened (O) and closed (C) periodically to ensure oxygen levels do not fall too low (short lines).

**Figure S2.**
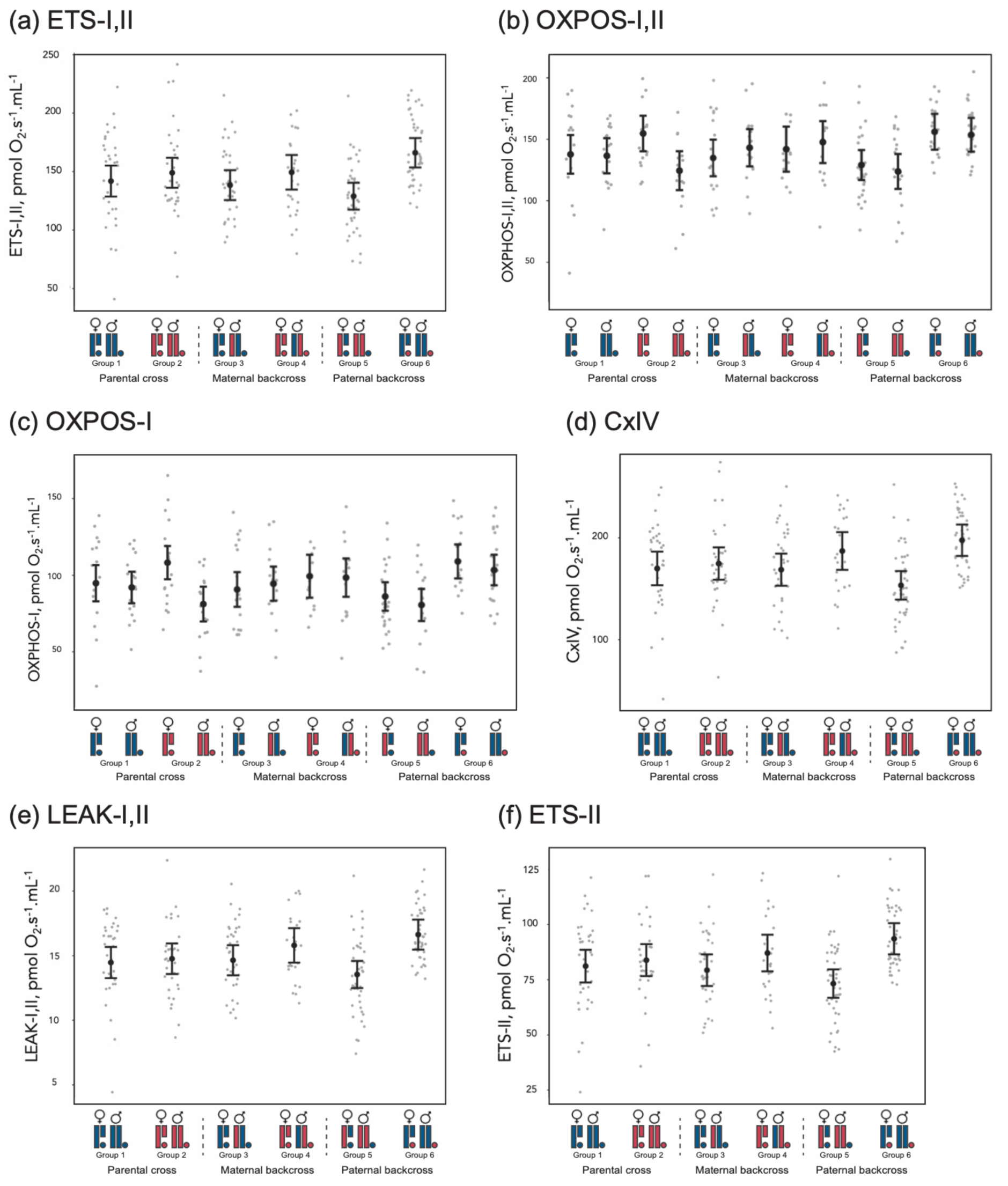
High resolution respiratory data from the six measures across the six experimental groups and split by sex when there was a significant interaction (Table 3). Datapoints indicate raw data per individual, error bars are 95% CIs based on model output.

**Figure S3.**
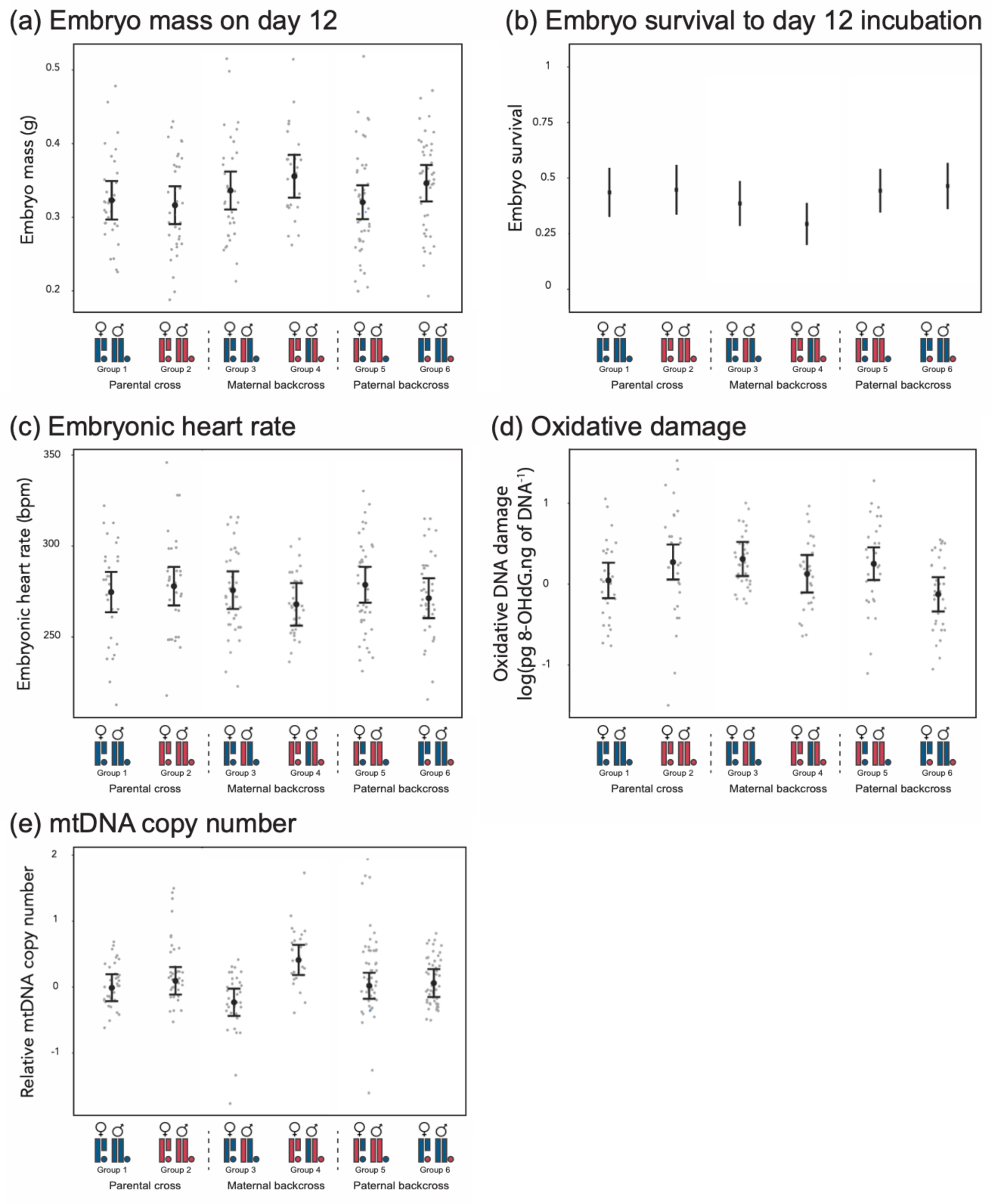
Non-respiratory measures made from embryo compared across the experimental groups, where each point is the raw data from an embryo and the error bars indicate the mean and 95% CI. A: embryo mass on day 12 of incubation, B: embryo survival to day 12 incubation (yes 1, or no 0), C: embryonic heart rate, calculated from a curve generated for each embryo, here shown the predicted heart rate 60 seconds after removal from the incubator, D: 8-OHdG oxidative DNA damage, E: mtDNA copy number relative to the nuDNA copy number.

